# Heatwaves strengthen historical contingency in species interactions: evidence from nectar microbes

**DOI:** 10.64898/2026.07.21.739338

**Authors:** Rosa M. McGuire, Cailinn M. C. Allen, Hlee Xiong, Nathan Vong, Tadashi Fukami

## Abstract

Heatwaves can affect species abundances by changing how species interact in local communities. However, these effects remain poorly understood, especially when the outcome of species interactions is contingent on their arrival history. We studied how heatwaves affect interactions between the yeast *Metschnikowia reukaufii* and the bacterium *Acinetobacter nectaris*, both found in the floral nectar of *Diplacus aurantiacus*, a hummingbird-pollinated shrub native to California. Microbes were introduced to artificial nectar in different orders of arrival in the presence or absence of simulated two-day heatwaves. Heatwaves made yeast–bacterium interactions more contingent on arrival history, causing large variation in nectar acidity, a factor known to affect hummingbird preference. Specifically, in the absence of heatwaves, *Acinetobacter* became abundant regardless of arrival history, suppressing *Metschnikowia* and reducing nectar pH. In contrast, in the presence of heatwaves, *Acinetobacter* dominance depended on arrival history and heatwave timing. If *Acinetobacter* arrived after *Metschnikowia* during a heatwave, it was suppressed by *Metschnikowia* substantially enough to keep nectar pH at a high level. These results suggest that heatwaves shift species interactions from determinism to historical contingency, with taxonomic and functional consequences.

## INTRODUCTION

As extreme climatic events become more common (Allan et al. 2023), increasing attention is being paid to understanding ecological consequences in various ecosystems (Vázquez et al. 2017; Maxwell et al. 2019; Cinto Mejía and Wetzel 2023). Studies have shown, for example, that heatwaves, or the extended periods of excessive heat (Perkins-Kirkpatrick and Lewis 2020; Bunting et al. 2024), influence species abundances not just directly by favoring heat-tolerant species over heat-sensitive ones, but also indirectly by changing the way species interact with one another in local communities (Chen and Lewis 2023; Barnes et al. 2024). Numerous studies have examined the effects of temperature on species interactions, but most of this work has focused on species performance at constant temperature (e.g., Lax et al. 2020; Lyberger et al. 2026). Heatwaves are characterized by fluctuating, not constant, temperatures, where the timing of events can be critical for understanding their consequences. Given this knowledge gap, one unresolved question is whether heatwaves make species interactions more deterministic or more historically contingent.

Deterministic interactions are those that yield the same outcome in terms of local species abundances regardless of the history of species arrival in community assembly (Drake 1991). Collectively, deterministic interactions should homogenize local communities in a region. In contrast, historically contingent interactions are those where the outcome of interactions depends on the order and timing of species arrival (Fukami 2015). These interactions can contribute to generating variation in species composition among local communities, enhancing taxonomic and functional diversity at a regional scale (Chase 2003; Wittmann and Fukami 2018). Theory and empirical evidence suggest that the extent of this historical contingency depends on environmental variability (Tucker and Fukami 2014; Hawkes and Keitt 2015; Grainger et al. 2018). However, environmental variability is logistically difficult to study (Hardison et al. 2026) and much remains unknown about how heatwaves may alter the extent of historical contingency. Addressing this question in a study system that is simple and easy to replicate and manipulate in a realistic context of heatwaves would help to advance knowledge on this topic.

This paper reports such a study. It focused on two species of nectar-specialist microorganisms, one yeast and one bacterium, commonly found in the floral nectar of the sticky monkeyflower, *Diplacus* (formerly *Mimulus*) *aurantiacus*. This hummingbird-pollinated shrub is native to California, where heatwaves are projected to increase in frequency, intensity, and duration (Gershunov and Guirguis 2012; Palipane and Grotjahn 2018). Previous work has shown that interactions between yeasts and bacteria in *D. aurantiacus* nectar can be historically contingent, depending on environmental conditions of the nectar habitat (Tucker and Fukami 2014; Chappell et al. 2022). If bacteria arrive first, they can grow rapidly and acidify nectar so greatly that later-arriving yeasts would not be able to grow. This bacterial dominance has consequences for plant reproduction: hummingbirds are less likely to visit flowers once nectar is acidified and seed set is reduced (Vannette et al. 2013). On the other hand, if yeasts arrive first, they can reduce the amount of nutrients available in nectar so extensively that later-arriving bacteria would not grow, thereby preventing bacteria-induced reduction in hummingbird visitation and seed production (Chappell et al. 2022). Although these possibilities have been documented, they have not been examined in the context of heatwaves.

We tested the hypothesis that heatwaves would make yeast–bacterium interactions more deterministic through manipulation of species arrival history under simulated heatwaves. Based on data on microbial thermal performance (**Fig. 1A**), we predicted that heatwaves would weaken historical contingency because both yeasts and bacteria would suffer from high temperatures, weakening species interactions and therefore historical contingency. Specifically, we expected that bacteria, with a wider tolerance range at the high-temperature extreme, would deterministically outcompete yeasts under heatwave conditions.

**Fig. 1.**
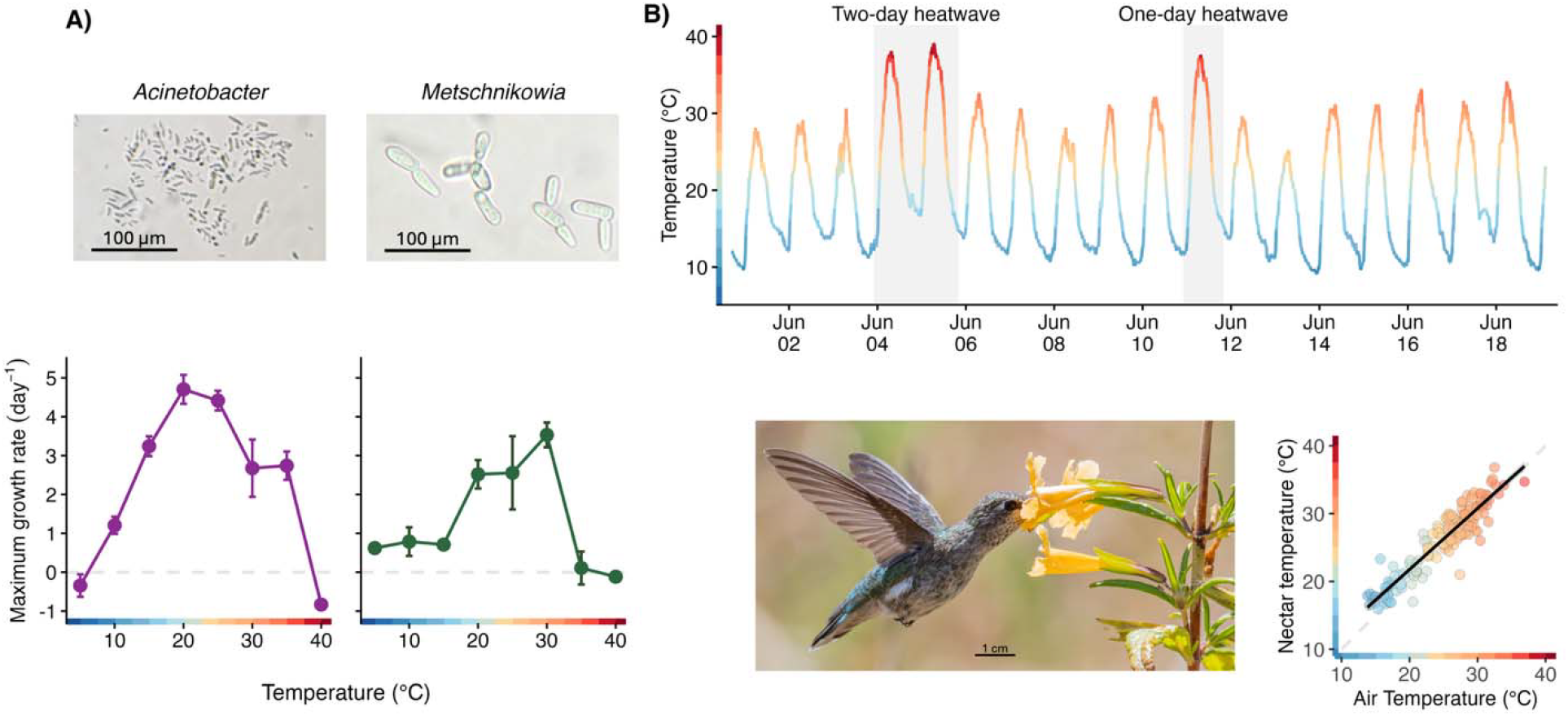
Study system. **A)** Maximum growth rate (mean +/-SE) of *Acinetobacter nectaris* and *Metschnikowia reukaufii* measured under different constant temperatures (images by Rosa McGuire). **B)** Ambient temperature recorded in June 2024 by a data logger (iButton) placed within a meter from a *D. aurantiacus* plant at Jasper Ridge Biological Preserve (‘Ootchamin ‘Ooyakma), showing two instances of heatwaves, indicated by shading. The temperature of floral nectar of *D. aurantiacus*, which is primarily pollinated by Anna’s Hummingbird (*Calypte anna*), was tightly coupled with air temperature (hummingbird image by Rick Morris). Points in the plot represent individual temperature measurements of floral nectar and of air (within 2 cm of the flower) for 49 *D. aurantiacus* flowers, with multiple measurements at different times of the day from each flower. Black line shows linear regression. Data collection and analyses are described in the Supplementary Material.

## METHODS

### Study organisms

We used the nectar-specialist bacterium *Acinetobacter nectaris* (strain SAP 949.2B; Álvarez-Pérez et al. 2021; hereafter *Acinetobacter*) and the nectar-specialist yeast *Metschnikowia reukaufii* (strain MR1; Peay et al. 2012; hereafter *Metschnikowia*), both isolated from the floral nectar of *D. aurantiacus* at Jasper Ridge Biological Preserve (‘Ootchamin ‘Ooyakma) (JRBP(‘O’O)), located in the Santa Cruz Mountains of California. These are some of the most commonly observed species of yeast and bacteria in *D. aurantiacus* nectar at JRBP(‘O’O) (Chappell et al. 2022; Belisle et al. 2012; Vannette and Fukami 2017). At this location, *D. aurantiacus* blooms from April to June, when ambient and nectar temperatures typically undergo daily fluctuations between 10 and 30 °C (**Fig. 1B**). During this period, heatwaves occur occasionally, with daily maximum temperatures reaching 40 °C and minimum temperatures remaining above 20 °C (**Fig. 1B**). The intensity, frequency, and duration of these heatwaves are projected to increase, with median duration doubling and hottest days warming by 3.5-6.0 °C over the next several decades (Palipane and Grotjahn 2018). Thermal performance curves show that these projected heatwaves negatively affect *Acinetobacter* and *Metschnikowia* (**Fig. 1A**). Their growth rate may start to decline at around 25-30 °C, reaching a negative value at 40 °C. However, it is not known how heatwaves affect the two species when they interact in the same flower.

### Inoculant preparation

*Acinetobacter* and *Metschnikowia* were streaked on tryptic soy agar (TSA) with 0.1 g/L of cycloheximide and yeast malt agar (YMA) with 0.1 g/L of chloramphenicol, respectively, from the glycerol stocks of the original isolates of the strains that were kept at -80 °C. The plates were incubated for two days at 25 °C. Four single colonies of each species were resuspended each in 3 mL of tryptic soy (TS) or yeast malt (YM) broth, respectively, and incubated for approximately 16 h at 26 °C and 200 rpm. These overnight cultures were distributed in 2-mL Eppendorf tubes and centrifuged at 6000 g for five minutes. The broth was then decanted, and 200 µL of artificial nectar added. Artificial nectar consisted of approximately 20% sucrose, 2% glucose, and 4% fructose (0.198, 0.024, and 0.043 g per mL, respectively) plus 3.16 mM amino acids from digested casein (OmniPur Casamino Acids, Millipore Sigma), following Vannette and Fukami (2014). Artificial nectar was filter-sterilized using a 0.2-µm filter before use. The centrifuging, decanting, and rinsing steps were repeated twice. Cell density in each culture was then determined by hemocytometer counts, and each culture diluted to 2000 cells per µL. These solutions were used to make replicated freezer stocks by mixing them into a glycerol solution to a final concentration of 15%, aliquoting them in many 200-µL PCR tubes, and storing the tubes at -80 °C so that the initial microbial densities would be identical across the repeated runs of the experiment (Francis et al. 2023). A control solution, consisting of artificial nectar and 15% glycerol, was also prepared and stored at -80 °C.

### Experimental design

We introduced *Acinetobacter* and *Metschnikowia* into artificial flowers alone (monocultures) or together (cocultures). Artificial flowers consisted of 200-µL PCR tubes, each containing 9 µL of artificial nectar in monocultures or 8 µL in cocultures. On each inoculation day, freezer stocks of *Acinetobacter, Metschnikowia*, and a control solution were thawed, vortexed, and diluted 10-fold in artificial nectar before addition to the artificial flowers.

For monocultures, 1 µL of the microbial stock was introduced to each of the PCR tubes. For cocultures, we used three introduction orders: *Acinetobacter* first, *Metschnikowia* first, and simultaneous. In the *Acinetobacter*-first and *Metschnikowia*-first treatments, we added 0.5 µL of the first species and 0.5 of the control solution on day 0, and 0.5 µL the second species and 0.5 of control solution on day 1. In the simultaneous introduction treatment, we added 0.5 µL of each species on day 0, and 1 µL of control solution on day 1. Thus, the final volume in all artificial flowers was 10 µL after species introduction was complete, an amount within the range of possibilities in real *D. aurantiacus* flowers (Peay et al. 2012). After each introduction, each microbial stock was diluted in 0.85% NaCl (10x), and 50 µL of each diluted stock was plated onto YM or TSA plates to determine inoculum density. Artificial flowers were destructively sampled daily over the five-day period, which was close to the average lifespan (about seven days) of individual *D. aurantiacus* flowers at JRBP (‘O’O) (Peay et al. 2012).

Each experimental run consisted of 100 artificial flowers (i.e., (2 monocultures + 3 cocultures) x 5 days x 4 replicates). We had four thermal regimes, with each conducted in a separate experimental run. One regime served as a control, in which temperature fluctuated daily between 10 and 25 °C (**Fig. 2A**), simulating typical conditions at JRBP (‘O’O) (**Fig. 1B**). The remaining three regimes included a two-day heatwave, with temperatures ranging from 20 °C to 40 °C, superimposed on the normal daily fluctuation between 10 and 25 °C. These regimes differed only in the timing of the heatwave relative to microbial introduction, as shown in **Fig. 2 B-D**. Artificial flowers were placed in an aluminum PCR tube block within an EchoTherm dry bath (Torrey Pines Scientific IC25) programmed at their respective thermal regime.

**Fig. 2.**
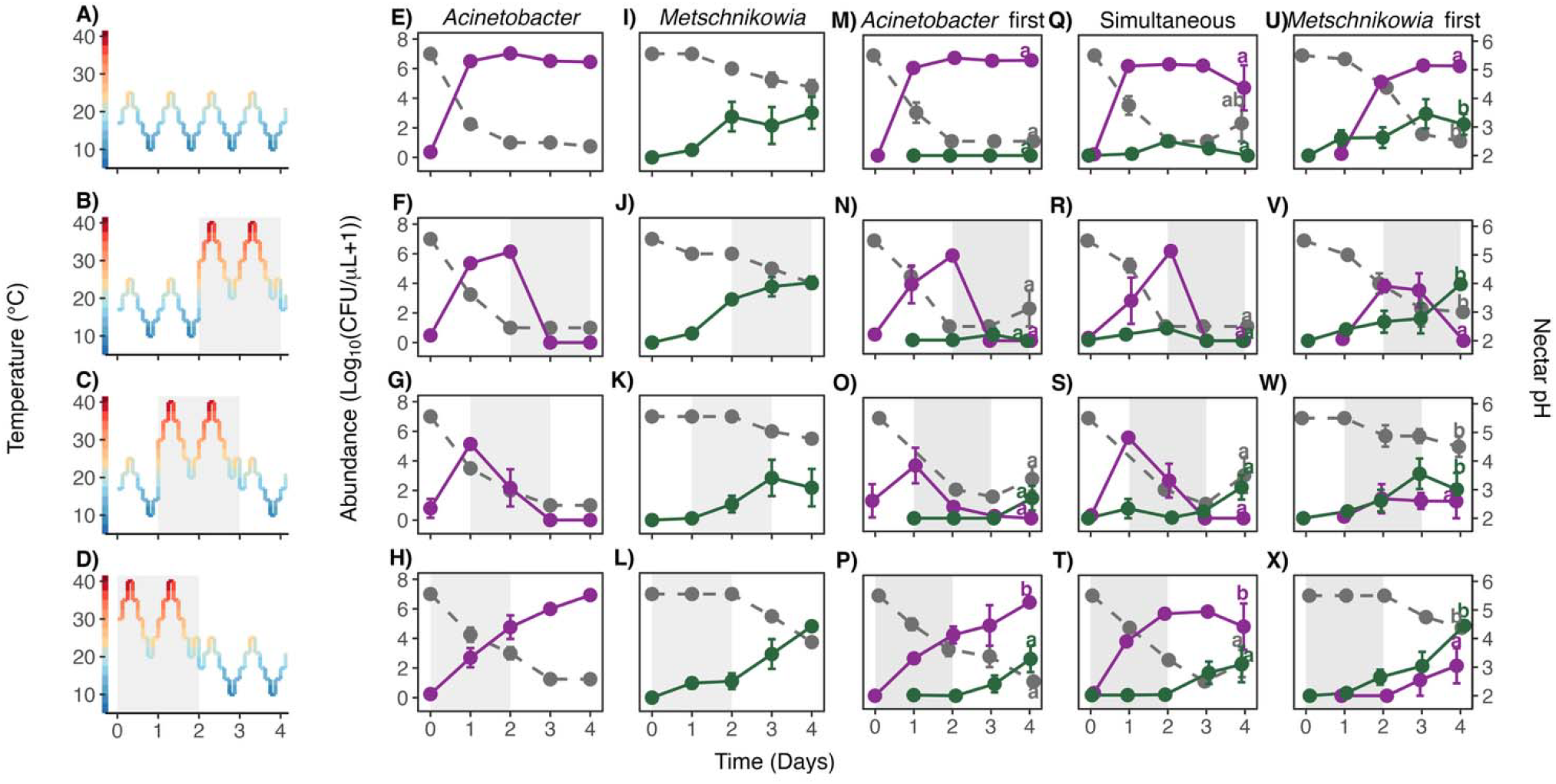
Heatwave effects. **A-D)** Temperature regimes used in the experiment, including a no-heatwave control (A), and two-day heatwaves that begin on days 2 (B), 1 (C), or 0 (D) of the experimental run. **E-X)** Response of *Acinetobacter nectaris* and *Metschnikowia reukaufii* to heatwaves in monocultures (E-L) and cocultures (M-X). Shading indicates time under heatwave conditions. Columns correspond to a different microbial inoculation treatment. Solid lines indicate microbial abundance, measured as log_10_(CFU/µL +1) (mean +/-SE). Grey dashed lines indicate nectar pH (mean +/-SE). See **Table S1** for statistical analysis and **Fig. S2** for plots with raw data. Letters indicate treatments that differed significantly (Tukey-adjusted comparisons).

### Data collection

In each experimental run, 20 artificial flowers (i.e., (2 monocultures + 3 cocultures) x 4 replicates) were destructively sampled on each of the five days. Artificial flowers were first vortexed for 3–5 seconds. To measure microbial abundance, we used five µL of nectar from each replicate for six serial dilutions in 45 µL of 0.85% NaCl, resulting in 10^-1^ to 10^-6^ dilutions. All dilutions were plated on 90 x 90 mm square plates (Simport Scientific D210-16) of TSA (supplemented with 0.1 g/L of cycloheximide) or YM (supplemented with 0.1 g/L of chloramphenicol). For cocultures, dilutions were plated onto both TSA and YM plates to determine bacterial and yeast abundance, respectively. Two 10-µL samples of each dilution were plated in each corresponding plate type. Plates were left to dry for 20 minutes and then placed in a 25 °C incubator. To measure nectar pH, two 0.5 µL drops of nectar were pipetted on a pH strip (range: 2.0–9.0, EMD Millipore 109584). After incubating for 3 days, microbial density was estimated by counting colony-forming units (CFU). For each artificial flower, 2-3 countable dilutions were recorded. Technical replicates (drops) were aggregated and averaged to estimate CFU per µL of nectar for each replicate on each day at each experimental condition.

### Data analysis

We used linear mixed models (LMMs) for monocultures and cocultures separately using the R package `lmerTest` (Kuznetsova et al. 2017) in R version 4.6.0 (R Core Team 2026). For monocultures, we used the abundance of yeast or bacteria (log_10_(CFU per µL + 1)) averaged across days 2 to 4 of the experiment as the response variable. We included thermal regime as fixed effect and experimental replicate as random effect. For cocultures, we also used the averaged abundance of yeast or bacteria on days 2-4 (log_10_(CFU per µL + 1)) as the response variable, and included thermal regime, arrival order, and their interaction as the explanatory variables, with experimental replicate as a random effect. A second set of LMMs was used to examine the effects of thermal regime and arrival order (if applicable) on pH for monocultures and cocultures, in the same way as described above. Treatment comparison was evaluated using ANOVA in the package `lmerTest` (Kuznetsova et al. 2017), and post-hoc pairwise comparisons (Tukey’s HSD) were conducted using the package `emmeans` (Lenth et al. 2019).

The entire experiment was repeated twice, with four replicates in each round. Despite large differences in initial inoculation density, the two rounds showed consistent patterns. Results from the first round are reported below while second round results can be found in the Supplementary Material (**Figs. S1, S3**).

## RESULTS

### Monocultures

In *Acinetobacter* monocultures, thermal regime significantly affected both microbial abundance (type II ANOVA, F_3,9_ = 179.5, p < 0.001) and nectar pH (type II ANOVA, F_3,9_ = 13.8, p = 0.0010, **Table S1, Fig. 2E-H**). Under the control thermal regime, *Acinetobacter* reached its maximum abundance within one day and maintained that density for the remainder of the experiment (**Fig. 2E**). This rapid population growth was accompanied by a decline in nectar pH from 5.5 to 2.5 by day 2 (**Fig. 2E**). When *Acinetobacter* arrived one or two days before the onset of the heatwave, the heatwave caused a rapid population decline, resulting in extirpation by day 3 (**Fig. 2F, G)**. However, nectar pH was kept at 2.5 even after the population crashed, and the final pH level was identical to the no-heatwave control (**Fig. 2F, G, Table S1A**). When *Acinetobacter* arrived the same time the heatwave started, growth was relatively slow during the heatwave, but the bacteria eventually reached the same maximum density as it did in the absence of the heatwave (**Fig. 2H**). Similarly, nectar pH declined more slowly but reached the same low level as in the no-heatwave control (**Fig. 2H**).

In *Metschnikowia* monocultures, thermal regime affected pH (type II ANOVA , F_3,9_ = 9.7742, p = 0.0034, **Table S1B**), but not overall abundance (type II ANOVA , F_3,9_ = 1.0489, p = 0.42, **Table S1B, Fig. 2I-L**). In the absence of the heatwave, *Metschnikowia* reached its maximum abundance by day 2 and reduced nectar pH to 4.5 by day 4 (**Fig. 2I**). When *Metschnikowia* arrived one or two days before heatwave onset, the heatwave hardly affected population growth or pH reduction (**Fig. 2J, K**), both behaving similarly to the no-heatwave control (**Table S1B**). When *Metschnikowia* arrived at the onset of the heatwave, it showed slower growth during the heatwave but reached a similar density to the maximum in the absence of the heatwave (**Fig. 2L**). Similarly, reduction in nectar pH was slower, but eventually led to the same low level, at 4-4.5 (**Fig. 2L**).

### Cocultures

When *Acinetobacter* and *Metschnikowia* were introduced together, population and pH outcomes depended on the interaction between arrival history and heatwave timing (**Table S2, Fig. 2M-X**). When *Acinetobacter* arrived before or simultaneously with *Metschnikowia* (**Fig. 2M-T**), it underwent abundance changes and reduced nectar pH as in monocultures (**Fig. 2E-H**) regardless of heatwave timing. In contrast, when *Metschnikowia* arrived first, it attained high abundance across all thermal regimes, and *Acinetobacter* abundance became contingent on heatwave timing (**Fig. 2U-X**). If *Acinetobacter* had at least one day to establish before the heatwave, it attained high abundance and reduced nectar pH to ≤ 3.0 (**Fig. 2U, V**). However, when introduced during the heatwave, *Acinetobacter* was strongly suppressed by *Metschnikowia*, resulting in nectar pH remaining near 4.5 (**Fig. 2W, X, Table S2C**).

## DISCUSSION

Contrary to our expectation, the results indicate that heatwaves shift the interactions between *Acinetobacter* and *Metschnikowia* in floral nectar from strong determinism to strong historical contingency in this system. Specifically, in the absence of heatwaves, *Acinetobacter* deterministically reached an abundance that was high enough to reduce nectar pH to 3 regardless of the timing of their arrival relative to *Metschnikowia* (**Fig. 2M, Q, U**). In doing so, *Acinetobacter* suppressed *Metschnikowia* because *Metschnikowia* could not grow well under low nectar pH (Tucker and Fukami 2014; Chappell et al. 2022). Some historical contingency was observed: *Metschnikowia* attained higher final abundance if it was introduced before, but not after, *Acinetobacter*. By and large, however, the *Acinetobacter*–*Metschnikowia* interaction and its consequence for nectar pH were deterministic. In contrast, when *Acinetobacter* arrived during a heatwave, its success depended strongly on arrival history (**Fig. 2P, T, X**). Early establishment of *Metschnikowia* suppressed *Acinetobacter* growth and maintained nectar pH near 4.5 (**Fig. 2X**), whereas when *Acinetobacter* arrived first, it dominated the community and reduced nectar pH to approximately 3 (**Fig. 2P, T**). Taken together, these findings suggest that heatwaves strengthened historical contingency of the otherwise largely deterministic *Acinetobacter*-*Metschnikowia* interactions, with strong effects on nectar pH.

Although the effects of heatwaves on species interactions have already been studied in other organisms (Ma et al. 2018; Buckley et al. 2025; Russell and McFrederick 2022), our study highlights the previously unrecognized possibility that heatwaves make arrival order matter to species abundances in local communities. Moreover, the variation in nectar pH that results from this historical contingency (**Fig. S4**) is known to have cascading effects on plant–pollinator mutualism (Chappell et al. 2022; Vannette et al. 2013; Rering et al. 2018; Martin et al. 2022), suggesting that historical contingency in yeast–bacterium interactions has both taxonomic and functional consequences. Previous work on *D. aurantiacus* has shown that nectar pH is not only a mechanism underlying interactions between *Acinetobacter* and *Metschnikowia*, but also a determinant of the functional consequences of nectar microbial community assembly. Because nectar pH affects hummingbird foraging preferences (Chappell et al. 2022; Vannette et al. 2013), *Acinetobacter*-driven reductions in pH decrease hummingbird visitation to *D. aurantiacus*, leading to reduced pollination and seed set (Chappell et al. 2022; Vannette et al. 2013). Thus, pH-mediated microbial interactions generate historical contingency that extends from microbial community assembly to plant–pollinator interactions. What is new about the results reported here is the possibility that heatwaves make this multi-level cascade contingent on the flower-scale history of microbial arrival. Since both *Acinetobacter* and *Metschnikowia* are also commonly found in many other plant species (Lachance et al. 2001; Vannette 2020), and because heatwaves are increasing in frequency and intensity in many parts of the world (Perkins-Kirkpatrick and Lewis 2020; Bunting et al. 2024), similar changes in yeast–bacterium interactions in nectar may be happening elsewhere as well.

Several caveats about our work point to future research directions. First, this study did not consider the physiological response of the host plants to heatwaves. Nectar quality and quantity may be affected by heatwaves, which can in turn affect yeast–bacterium interactions. Second, although hummingbirds are the primary pollinators of *D. aurantiacus* at our study site, insects also visit the flowers (Belisle et al. 2012; Vannette and Fukami 2017). Hummingbirds’ feeding behavior may be insensitive to temperature (Lawrence and Hazlehurst 2023), but insect visits to flowers are likely affected by heatwaves (Hemberger et al. 2023), resulting in changes in microbial dispersal across flowers. The effects of heatwaves on vector-mediated dispersal remain an unexplored question. Third, this study used only one strain per species, ignoring the role of strain diversity in microbial persistence in the field (Hughes et al. 2008). The outcome of species interactions and nectar chemistry might have differed with different strains of *Acinetobacter* and *Metschnikowia* (Chappell et al. 2022). Finally, many flowers continue to produce nectar, replenishing the nutrients available to the microbes, which we did not consider here.

In conclusion, this study has provided experimental evidence suggesting that extreme weather events strengthen historical contingency in local species interactions, which can in turn increase variability in species dominance and functioning among local communities. In our case, heatwaves appeared to benefit *Metschnikowia* by giving it the opportunity to competitively exclude *Acinetobacter* if it arrives first. An overall increase in *Metschnikowia* abundance could help to strengthen plant–pollinator mutualism by allowing nectar pH and therefore pollinator preference to be kept high when flowers have specific arrival history of nectar microbes. The extent to which this possibility is generalizable or specific to our study system remains to be investigated. We suggest that a deeper understanding of the effects of extreme climatic events on species interactions can be gained by considering how they modulate historical contingency.

## ACKNOWLEDGMENTS

We thank members of the Community Ecology Group at Stanford University for comments and Julio Meza for their assistance with preliminary experiments. H.X. was supported by the Stanford bioBUDS program. N.V. was supported by the Community College Outreach Program at Stanford. This research was supported by a National Science Foundation Postdoctoral Research Fellowship in Biology (DBI-2305992) awarded to R.M.M.

## REFERENCES

Allan, R. P., P. A. Arias, S. Berger, et al. 2023. “Intergovernmental Panel on Climate Change (IPCC). Summary for Policymakers.” In Climate Change 2021: The Physical Science Basis. Contribution of Working Group I to the Sixth Assessment Report of the Intergovernmental Panel on Climate Change. Cambridge University Press.

Álvarez-Pérez, S., L. J. Baker, M. M. Morris, et al. 2021. “Acinetobacter pollinis Sp. Nov., Acinetobacter baretiae Sp. Nov. and Acinetobacter rathckeae Sp. Nov., Isolated from Floral Nectar and Honey Bees.” International Journal of Systematic and Evolutionary Microbiology 71 (5): 004783.

Barnes, A D., J. R. Deslippe, A. M. Potapov, A. L. Romero-Olivares, L. A. Schipper, and C. J. Alster. 2024. “Does Warming Erode Network Stability and Ecosystem Multifunctionality?” Trends in Ecology & Evolution 39 (10): 892–94.

Belisle, M., K. G. Peay, and T. Fukami. 2012. “Flowers as Islands: Spatial Distribution of Nectar-Inhabiting Microfungi among Plants of Mimulus aurantiacus, a Hummingbird-Pollinated Shrub.” Microbial Ecology 63 (4): 711–18.

Buckley, L. B., R. B. Huey, and C.-S. Ma. 2025. “How Damage, Recovery, and Repair Alter the Fitness Impacts of Thermal Stress.” Integrative and Comparative Biology 65 (4): 1061–75.

Bunting, E. L., V. Tolmanov, and D. Keellings. 2024. “What Is a Heat Wave: A Survey and Literature Synthesis of Heat Wave Definitions across the United States.” PLOS Climate 3 (9): e0000468.

Chappell, C. R., M. K. Dhami, M. C. Bitter, et al. 2022. “Wide-Ranging Consequences of Priority Effects Governed by an Overarching Factor.” eLife 11 (October): e79647.

Chase, J. M. 2003. “Community Assembly: When Should History Matter?” Oecologia 136 (4): 489–98.

Chen, J., and O. T. Lewis. 2023. “Experimental Heatwaves Facilitate Invasion and Alter Species Interactions and Composition in a Tropical Host-Parasitoid Community.” Global Change Biology 29 (22): 6261–75.

Cinto Mejía, E., and W. C. Wetzel. 2023. “The Ecological Consequences of the Timing of Extreme Climate Events.” Ecology and Evolution 13 (1): e9661.

Drake, J. A. 1991. “Community-Assembly Mechanics and the Structure of an Experimental Species Ensemble.” The American Naturalist 137 (1): 1–26.

Francis, J. S., T. G. Mueller, and R. L. Vannette. 2023. “Intraspecific Variation in Realized Dispersal Probability and Host Quality Shape Nectar Microbiomes.” New Phytologist 240 (3): 1233–45.

Fukami, T. 2015. “Historical Contingency in Community Assembly: Integrating Niches, Species Pools, and Priority Effects.” Annual Review of Ecology, Evolution, and Systematics 46: 1–23.

Gershunov, A., and K. Guirguis. 2012. “California Heat Waves in the Present and Future.” Geophysical Research Letters 39 (18).

Grainger, T. N., A. I. Rego, and B. Gilbert. 2018. “Temperature-Dependent Species Interactions Shape Priority Effects and the Persistence of Unequal Competitors.” The American Naturalist 191 (2): 197–209.

Hardison, E. A., E. Donham, M. Massey, A. J. Morash, and G. D. Schwieterman. 2026. “Incorporating Environmental Variability into Physiological Experiments: Rationale and Resources.” Philosophical Transactions of the Royal Society B: Biological Sciences 381 (1946): 20250053.

Hawkes, C. V., and T. H. Keitt. 2015. “Resilience vs. Historical Contingency in Microbial Responses to Environmental Change.” Ecology Letters 18 (7): 612–25.

Hemberger, J. A., N. M. Rosenberger, and N. M. Williams. 2023. “Experimental Heatwaves Disrupt Bumblebee Foraging through Direct Heat Effects and Reduced Nectar Production.” Functional Ecology 37 (3): 591–601.

Hughes, A. R., B. D. Inouye, M. T. J. Johnson, N. Underwood, and M. Vellend. 2008. “Ecological Consequences of Genetic Diversity.” Ecology Letters 11 (6): 609–23.

Kuznetsova, A., P. B. Brockhoff, and R. H. B. Christensen. 2017. “lmerTest Package: Tests in Linear Mixed Effects Models.” Journal of Statistical Software 82 (December): 1–26.

Lachance, M.-A., W. T. Starmer, C. A. Rosa, J. M. Bowles, J. S. F. Barker, and D. H. Janzen. 2001. “Biogeography of the Yeasts of Ephemeral Flowers and Their Insects.” FEMS Yeast Research 1 (1): 1–8.

Lawrence, S. L., and J. Hazlehurst. 2023. “Hummingbird Foraging Preferences during Extreme Heat Events.” Ecology and Evolution 13 (5): e10053.

Lax, S., C. I. Abreu, and J. Gore. 2020. “Higher Temperatures Generically Favour Slower-Growing Bacterial Species in Multispecies Communities.” Nature Ecology & Evolution 4 (4): 560–67.

Lenth, R., H. Singmann, J. Love, P. Buerkner, and M. Herve. 2019. “Package ‘Emmeans.’” R Package Version 1 (3.2): 1199.

Lyberger, K. P., J. Farner, E. A. Mordecai, and T. Fukami. 2026. “Warming Strengthens Priority Effects Driven by an Ontogenetic Shift from Host to Predator.” The American Naturalist, August 3, 743450.

Ma, C.-S., L. Wang, W. Zhang, and V. H. W. Rudolf. 2018. “Resolving Biological Impacts of Multiple Heat Waves: Interaction of Hot and Recovery Days.” Oikos 127 (4): 622–33.

Martin, V. N., R. N. Schaeffer, and T. Fukami. 2022. “Potential Effects of Nectar Microbes on Pollinator Health.” Philosophical Transactions of the Royal Society B: Biological Sciences 377 (1853): 20210155.

Maxwell, S. L., N. Butt, M. Maron, et al. 2019. “Conservation Implications of Ecological Responses to Extreme Weather and Climate Events.” Diversity and Distributions 25 (4): 613–25.

Palipane, E., and R. Grotjahn. 2018. “Future Projections of the Large-Scale Meteorology Associated with California Heat Waves in CMIP5 Models.” Journal of Geophysical Research: Atmospheres 123 (16): 8500–8517.

Peay, K. G., M. Belisle, and T. Fukami. 2012. “Phylogenetic Relatedness Predicts Priority Effects in Nectar Yeast Communities.” Proc Biol Sci 279 (1729): 749–58.

Perkins-Kirkpatrick, S. E., and S. C. Lewis. 2020. “Increasing Trends in Regional Heatwaves.” Nature Communications 11 (1): 3357.

R Core Team. 2026. R: A Language and Environment for Statistical Computing. R Foundation for Statistical Computing, Vienna, Austria. 10.32614/R.manuals.

Rering, C. C., J. J. Beck, G. W. Hall, M. M. McCartney, and R. L. Vannette. 2018. “Nectar-Inhabiting Microorganisms Influence Nectar Volatile Composition and Attractiveness to a Generalist Pollinator.” New Phytologist 220 (3): 750–59.

Russell, K. A., and Q. S. McFrederick. 2022. “Floral Nectar Microbial Communities Exhibit Seasonal Shifts Associated with Extreme Heat: Potential Implications for Climate Change and Plant-Pollinator Interactions.” Front Microbiol 13: 931291.

Tucker, C. M., and T. Fukami. 2014. “Environmental Variability Counteracts Priority Effects to Facilitate Species Coexistence: Evidence from Nectar Microbes.” Proc Biol Sci 281 (1778): 20132637.

Vannette, R. L. 2020. “The Floral Microbiome: Plant, Pollinator, and Microbial Perspectives.” Annual Review of Ecology, Evolution, and Systematics 51 (Volume 51, 2020): 363–86.

Vannette, R. L., and T. Fukami. 2014. “Historical Contingency in Species Interactions: Towards Niche□based Predictions.” Ecology Letters 17 (1): 115–24.

Vannette, R. L., and T. Fukami. 2017. “Dispersal Enhances Beta Diversity in Nectar Microbes.” Ecology Letters 20 (7): 901–10.

Vannette, R. L., M.-P. L. Gauthier, and T. Fukami. 2013. “Nectar Bacteria, but Not Yeast, Weaken a Plant–Pollinator Mutualism.” Proceedings of the Royal Society B: Biological Sciences 280 (1752): 20122601.

Vázquez, D. P., E. Gianoli, W. F. Morris, and F. Bozinovic. 2017. “Ecological and Evolutionary Impacts of Changing Climatic Variability.” Biological Reviews 92 (1): 22–42.

Wittmann, M. J., and T. Fukami. 2018. “Eco-Evolutionary Buffering: Rapid Evolution Facilitates Regional Species Coexistence despite Local Priority Effects.” The American Naturalist 191 (6): E171–84.

