## Supplementary Information for "Heatwaves strengthen historical contingency in species interactions: evidence from nectar microbes"

**Appendix S1: Supplementary material**

*Field measurements*

To understand the relationship between air and nectar temperature, we measured the internal temperature of 49 flowers over the course of six days in June and July of 2024 at the Stanford Dish trail (37°24'36.30"N 122°9'40.76"W). Temperature was measured using a flexible microprobe (IT-23, World Precision Instruments) connected to a thermocouple thermometer (Digi-Sense Temp100, WD-91428-03) in a way similar to Herrera and Medrano (2017). First, air temperature was measured and then the probe was inserted into the flower. Each flower was measured three or four times at different times of the day over the course of the experiment. Measurements were taken during early, mid-morning, and early afternoon to include a wide range of air temperatures. We fitted a linear mixed-effects model with nectar temperature as the response variable, air temperature as the explanatory variable, and flower ID as a random intercept using the R package lmerTest (Kuznetsova et al. 2017). R^2^ was calculated using the package performance (Lüdecke et al. 2021). Air temperature was a significant predictor of nectar temperature (p<0.001, marginal R^2^= 0.866, conditional R^2^= 0.89, **Fig. 1B**).

*Effects of temperature on growth rate*

Monocultures of *Acinetobacter nectaris* and *Metschnikowia reukaufii* prepared the same way as described in the methods section were replicated four times and incubated for five days at eight different constant temperatures, from 5 to 40 °C, in increments of 5 °C. Microbial abundance and nectar pH were measured daily for five days. For each measurement, six serial dilutions (10^-1^ to 10^-6^) were plated in triplicate, with each sample consisting of 10 µL, on square TSA (supplemented with 0.1 g/L of cycloheximide) or YM (supplemented with 0.1 g/L of chloramphenicol) plates. Colonies were counted after incubating for three days at 25 °C and 2-3 countable dilutions were recorded for each artificial flower. Technical replicates (drops) were aggregated and averaged to estimate CFU per µL of nectar for each replicate on each day at each experimental temperature. We used CFU data to estimate the maximum growth rate calculated as the maximum slope between days 0-2 of the experiment. Maximum growth rates were calculated for each replicate at each experimental temperature and used to build the temperature response curves shown in **Fig. 1A**.

**References**

1. Herrera, Carlos M., and Mónica Medrano. 2017. “Pollination Consequences of Simulated Intrafloral Microbial Warming in an Early-Blooming Herb.” *Flora*, Patterns and mechanisms in plant-pollinator interactions, vol. 232 (July): 142–49. https://doi.org/10.1016/j.flora.2016.10.003.
2. Kuznetsova, Alexandra, Per B. Brockhoff, and Rune H. B. Christensen. 2017. “lmerTest Package: Tests in Linear Mixed Effects Models.” *Journal of Statistical Software* 82 (December): 1–26. https://doi.org/10.18637/jss.v082.i13.
3. Lüdecke, Daniel, Mattan S. Ben-Shachar, Indrajeet Patil, Philip Waggoner, and Dominique Makowski. 2021. “Performance: An R Package for Assessment, Comparison and Testing of Statistical Models.” *Journal of Open Source Software* 6 (60): 3139. https://doi.org/10.21105/joss.03139.

**Appendix S2: Supplementary figures**

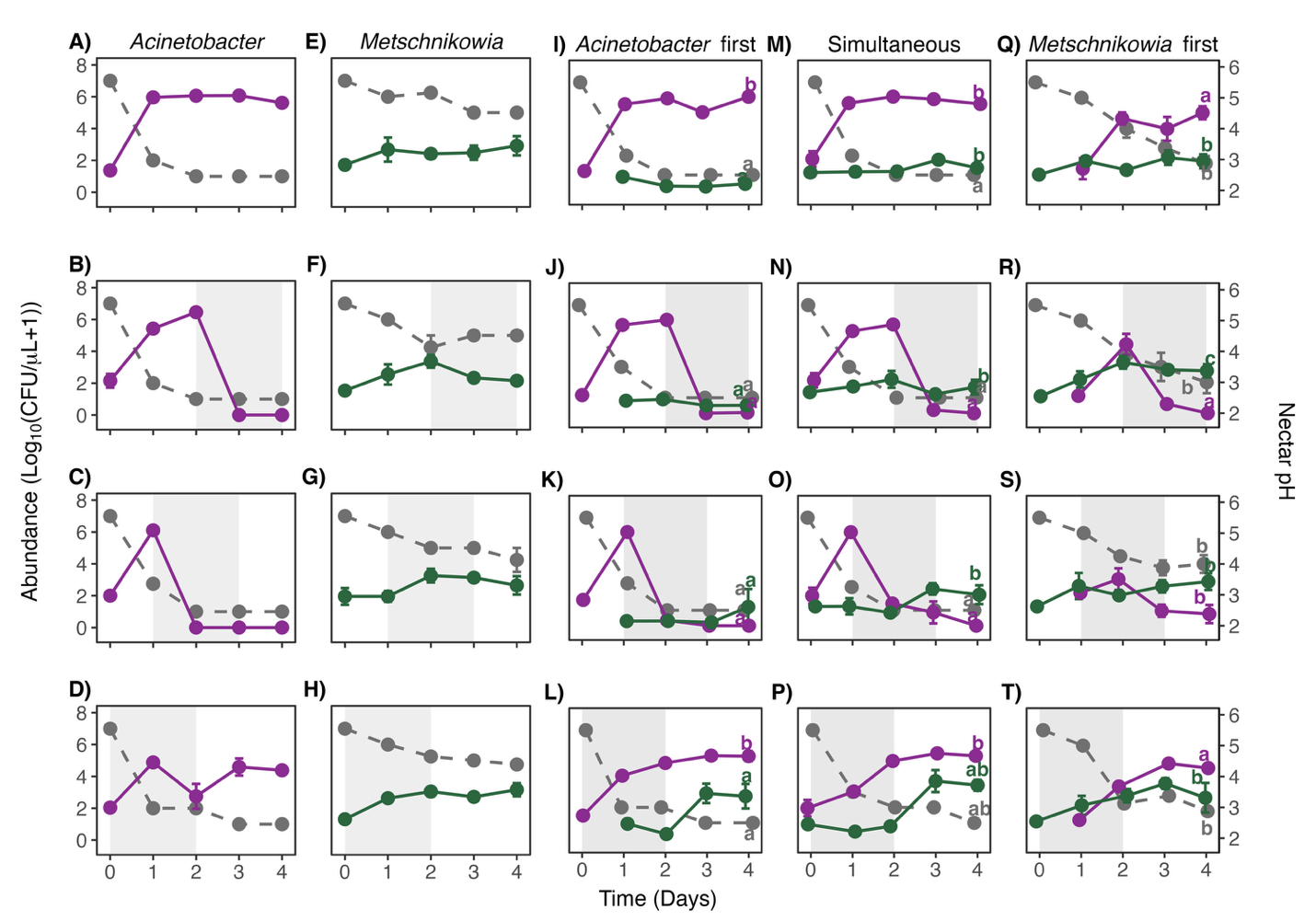

**Fig. S1.** Response of *Acinetobacter nectaris* and *Metschnikowia reukaufii* to heatwaves in monocultures (A-H) and cocultures (I-T) in the second experimental run. Shading, symbols, and letters are as in **Fig. 2**. See **Tables S3 and S4** for statistical analysis and **Fig. S3** for plots with raw data.

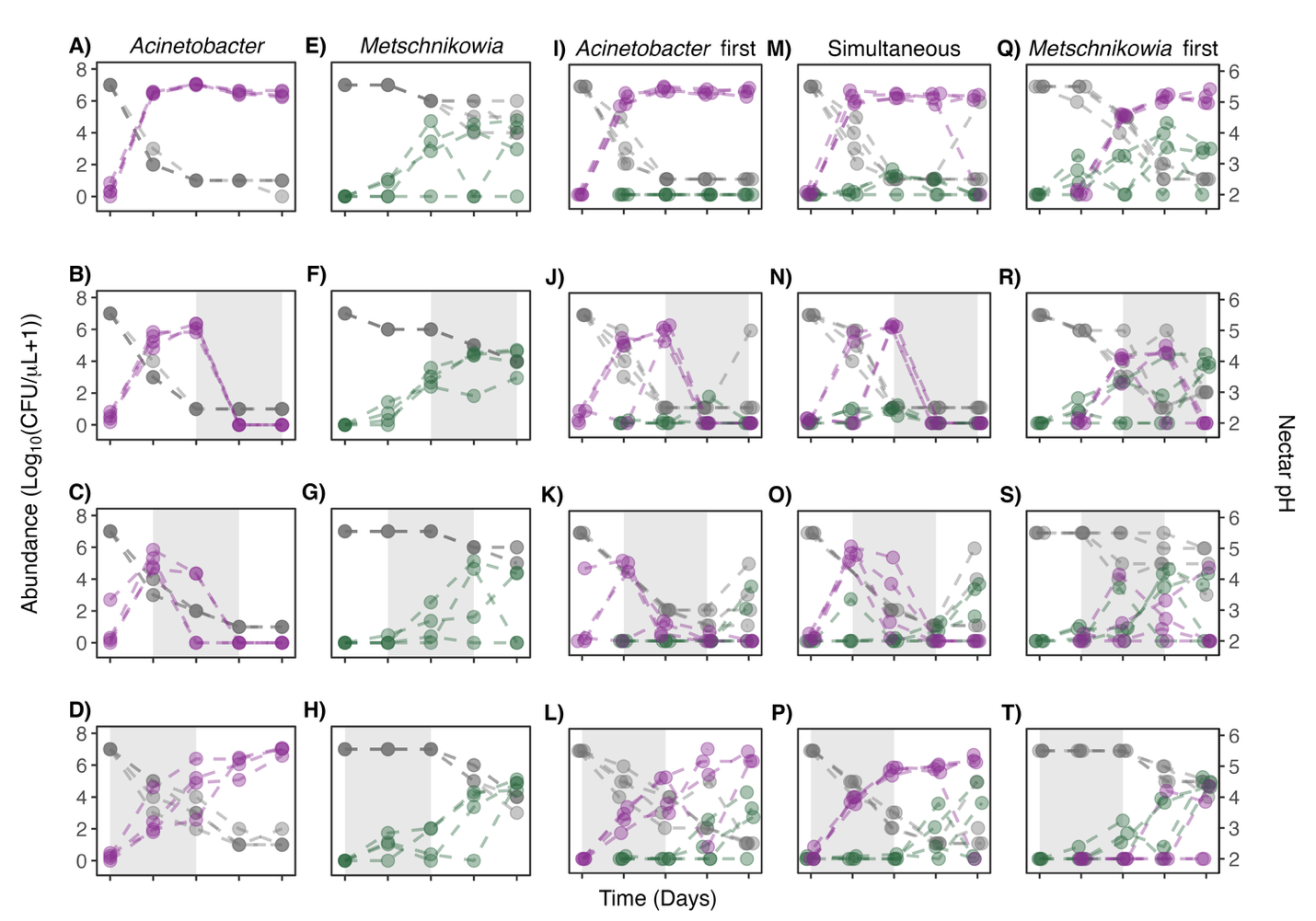

**Fig. S2.** Response of *Acinetobacter nectaris* and *Metschnikowia reukaufii* to heatwaves in monocultures (A-H) and cocultures (I-T) in the first experimental run. Shading, symbols, and letters are as in **Fig. 2**, but here raw data from each replicate are shown separately.

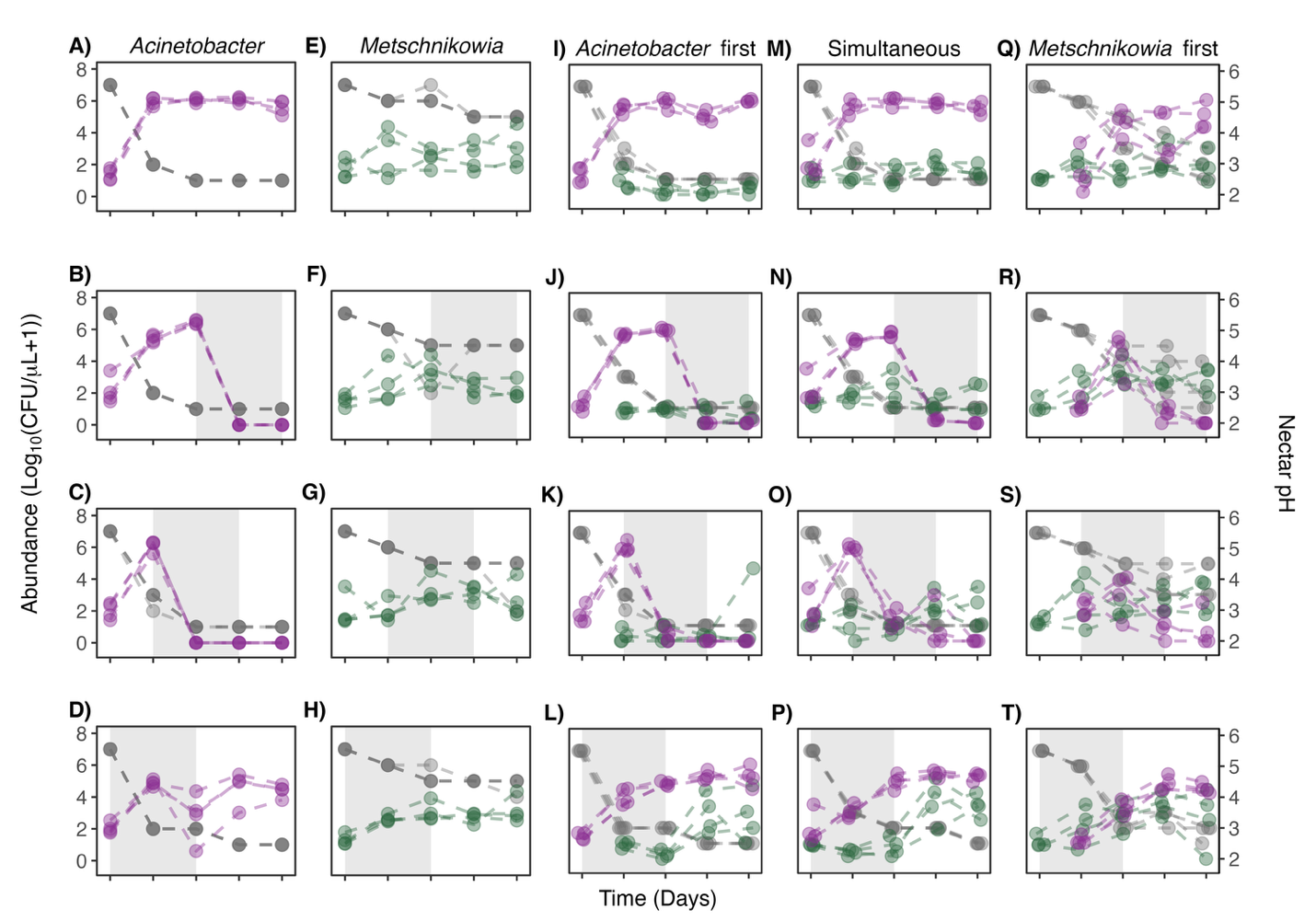

**Fig. S3.** Response of *Acinetobacter nectaris* and *Metschnikowia reukaufii* to heatwaves in monocultures (A-H) and cocultures (I-T) in the second experimental run. Shading, symbols, and letters are as in **Fig. S1**, but here raw data from each replicate are shown separately.

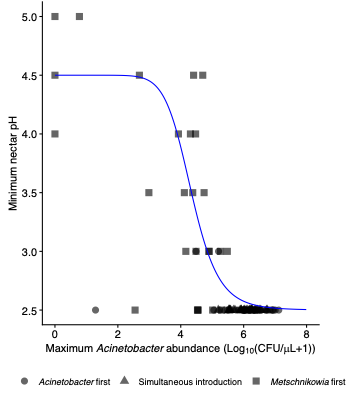

**Fig. S4.** Relationship between maximum *Acinetobacter* abundance after introduction, quantified as Log_10_(CFU/µL +1), and minimum nectar pH. Maximum and minimum values were obtained for each replicate and treatment for each experimental run. Shapes correspond to different arrival order treatments. Points correspond to experimental measurements across the two rounds of the experiment and blue line shows a fit to the Hill equation.

**Appendix S3: Supplementary tables**

**Table S1.** Results from a two-way ANOVA for the abundance (log_10_(CFU/µl +1)) and pH of A) *Acinetobacter* and B) *Metschnikowia* in monocultures and their relationship with temperature regime in the first experimental run. Text in bold indicates p-values < 0.05.

A) *Acinetobacter*

|  | Abundance | | | | pH | | | |
| --- | --- | --- | --- | --- | --- | --- | --- | --- |
| Variable | Num df | Den df | F | P | Num df | Den df | F | p |
| Temperature regime | 3 | 9 | 179.5 | **<0.001** | 3 | 9 | 13.8 | **0.0010** |

B) *Metschnikowia*

|  | Abundance | | | | pH | | | |
| --- | --- | --- | --- | --- | --- | --- | --- | --- |
| Variable | Num df | Den df | F | P | Num df | Den df | F | p |
| Temperature regime | 3 | 9 | 1.0 | 0.4173 | 3 | 9 | 9.8 | **0.0035** |

**Table S2.** Results from a two-way ANOVA for the abundance (log_10_(CFU/µl +1)) of A) *Acinetobacter* and B) *Metschnikowia* and C) pH in cocultures and their relationship with temperature regime, arrival order, or their interaction in the first experimental run. Text in bold indicates p-values < 0.05.

A) *Acinetobacter*

|  | Abundance | | | |
| --- | --- | --- | --- | --- |
| Variable | Num df | Den df | F | P |
| Temperature regime | 3 | 36 | 100.3 | **<0.001** |
| Arrival order | 2 | 36 | 6.5 | **0.0383** |
| Temperature x arrival order | 6 | 36 | 11.6 | **<0.001** |

B) *Metschnikowia*

|  | Abundance | | | |
| --- | --- | --- | --- | --- |
| Variable | Num df | Den df | F | P |
| Temperature regime | 3 | 36 | 4.2 | **0.0116** |
| Arrival order | 2 | 36 | 36.7 | **<0.001** |
| Temperature x arrival order | 6 | 36 | 0.3 | 0.9212 |

C) pH

|  | pH | | | |
| --- | --- | --- | --- | --- |
| Variable | Num df | Den df | F | P |
| Temperature regime | 3 | 33 | 22.1 | **<0.001** |
| Arrival order | 2 | 33 | 69.7 | **<0.001** |
| Temperature x arrival order | 6 | 33 | 4.9 | **0.0011** |

**Table S3.** Results from a two-way ANOVA for the abundance (log_10_(CFU/µl +1)) and pH of A) *Acinetobacter* and B) *Metschnikowia* in monocultures and their relationship with temperature regime in the second experimental run. Text in bold indicates p-values < 0.05. The model for pH in *Acinetobacter* monocultures was not fitted, because all values were identical (pH=2.5 across all treatments).

A) *Acinetobacter*

|  | Abundance | | | | pH | | | |
| --- | --- | --- | --- | --- | --- | --- | --- | --- |
| Variable | Num df | Den df | F | P | Num df | Den df | F | p |
| Temperature regime | 3 | 9 | 108.2 | **<0.001** | NA | NA | NA | NA |

B) *Metschnikowia*

|  | Abundance | | | | pH | | | |
| --- | --- | --- | --- | --- | --- | --- | --- | --- |
| Variable | Num df | Den df | F | P | Num df | Den df | F | p |
| Temperature regime | 3 | 9 | 1.0 | 0.4333 | 3 | 9 | 3.8 | 0.0538 |

**Table S4.** Results from a two-way ANOVA for the abundance (Log_10_(CFU/µl +1)) of A) *Acinetobacter* and B) *Metschnikowia* and C) pH in coculture and their relationship with temperature regime, arrival order, or their interaction in the second set of replicates. Text in bold indicates p-values < 0.05.

A) *Acinetobacter*

|  | Abundance | | | |
| --- | --- | --- | --- | --- |
| Variable | Num df | Den df | F | p |
| Temperature regime | 3 | 33 | 309.7 | **<0.001** |
| Arrival order | 2 | 33 | 4.3 | **0.0218** |
| Temperature x arrival order | 6 | 33 | 8.2 | **<0.001** |

B) *Metschnikowia*

|  | Abundance | | | |
| --- | --- | --- | --- | --- |
| Variable | Num df | Den df | F | p |
| Temperature regime | 3 | 33 | 13.1 | **<0.001** |
| Arrival order | 2 | 33 | 40.8 | **<0.001** |
| Temperature x arrival order | 6 | 33 | 1.4 | 0.2361 |

C) pH

|  | pH | | | |
| --- | --- | --- | --- | --- |
| Variable | Num df | Den df | F | p |
| Temperature regime | 3 | 33 | 2.1 | 0.1127 |
| Arrival order | 2 | 33 | 95.0 | **<0.001** |
| Temperature x arrival order | 6 | 33 | 6.13 | **<0.001** |
